# Coumarins link rhizobacteria perception in roots to systemic resistance in leaves

**DOI:** 10.64898/2026.08.11.741996

**Authors:** Shu-Hua Hsu, Max J.J. Stassen, Kevin Robe, Esther Izquierdo, Christian Dubos, Corné M.J. Pieterse, Ioannis A. Stringlis

## Abstract

Induced systemic resistance (ISR) is activated in leaves upon root colonization by beneficial microbes, yet the signals linking rhizosphere perception to shoot immunity remain unknown. In the *Arabidopsis thaliana-Pseudomonas simiae* WCS417 model interaction, the root-specific transcription factor MYB72 and its target gene *BGLU42* regulate ISR and the production, activation, and root secretion of coumarins, specialized metabolites involved in plant iron (Fe) acquisition and rhizosphere microbiome assembly. Overexpression of *BGLU42* confers constitutive ISR in leaves, suggesting a link between coumarin metabolism and systemic immunity. Here, two-photon multispectral imaging and targeted metabolite profiling revealed that, under Fe-sufficient conditions, WCS417 induces a distinct spatial pattern of F6’H1-dependent coumarin accumulation along the root system. These WCS417-induced coumarin signatures differed from those observed under Fe deficiency, indicating activation of a microbiota-specific coumarin metabolic program. Increased coumarin accumulation in roots was followed by a rise in coumarin levels in shoots. Time-resolved transcriptome profiling supported this metabolic reprogramming, showing rapid activation of Fe acquisition and coumarin biosynthesis genes in roots, including *F6’H1, MYB72,* and *BGLU42*, followed by delayed but similar transcriptional responses in shoots. Functional analyses demonstrated that coumarin biosynthesis is required for WCS417-ISR: the *f6’h1* mutant failed to mount systemic resistance, whereas *F6’H1* overexpression conferred constitutive resistance to bacterial and fungal pathogens. In addition, WCS417-mediated coumarin accumulation systemically modulated flg22-triggered reactive oxygen species production in leaves in an F6’H1-dependent manner. Together, our results identify coumarins as key mediators linking rhizobacterial perception in roots to systemic immune signaling and resistance in leaves.

## INTRODUCTION

Plant phenotypes are strongly shaped by interactions with the diverse microbial communities that inhabit the rhizosphere (Berendsen et al., 2012; Rolfe et al., 2019). Root exudates selectively recruit and structure these microbial assemblages, typically reducing microbial diversity while enriching for specific taxa, a process known as the “rhizosphere effect” (Hiltner, 1904; Poppeliers et al., 2024; Uribe Acosta et al., 2025). While some root-associated microbes can negatively impact plant performance through infection or competition for nutrients, many are beneficial, enhancing plant nutrition, suppressing pathogens, or activating plant immune responses. These intimate interactions with beneficial microbes have led to the emerging concept of the extended plant immune system (Pieterse, 2025), in which plants rely not only on intrinsic defense mechanisms but also on beneficial microbe-mediated protection to resist biotic stress. Within this framework, plant-associated microbiota are considered integral components of the plant’s immune capacity, capable of enhancing host resistance through immune-modulatory functions. One of the best-characterized manifestations of this extended immune system is induced systemic resistance (ISR), a state of enhanced defensive capacity in aboveground tissues triggered by specific root-associated beneficial microbes (Pieterse et al., 2014).

The root colonizing rhizobacterium *Pseudomonas simiae* WCS417 (WCS417) is a well-established model for beneficial plant-microbe interactions (Pieterse et al., 2021). In *Arabidopsis thaliana* (Arabidopsis), colonization of the roots by WCS417 promotes root architectural changes and plant growth (Zamioudis et al., 2013), and induces a broad-spectrum disease resistance in foliar tissues, termed WCS417-ISR (Pieterse *et al*., 2014). In Arabidopsis, WCS417-ISR depends on jasmonic acid (JA), ethylene (ET), and the defense regulator NPR1 (Pieterse et al., 1998). In roots, WCS417-ISR requires activation of the root-specific transcription factor MYB72 (Van der Ent et al., 2008), whose downstream target gene *β-GLUCOSIDASE42* (*BGLU42*) is essential for the ISR phenotype, as overexpression of *BGLU42* results in constitutive systemic resistance against multiple pathogens (Zamioudis et al., 2014).

In roots, MYB72 and BGLU42 also regulate the biosynthesis and activation of coumarins, specialized metabolites that contribute to iron (Fe) acquisition from the soil environment and, due to their selective antimicrobial activity, shape microbiome assembly in the rhizosphere (Harbort et al., 2020; Palmer et al., 2013; Robe et al., 2021a; Robe et al., 2025; Stringlis et al., 2019; Stringlis et al., 2018b; Voges et al., 2019; Zamioudis *et al*., 2014). Notably, WCS417 itself is largely insensitive to the antimicrobial effects of these coumarins, potentially allowing it to persist and even benefit from coumarin-mediated re-shaping of the root microbiome (Stringlis *et al*., 2018b). This suggests a feed-forward loop in which WCS417-induced activation of MYB72 and BGLU42 enhances coumarin biosynthesis and secretion, thereby promoting a rhizosphere environment that favors WCS417 colonization and reinforcement of ISR activation.

Beyond their more recently discovered role in nutrient acquisition and microbiome structuring, coumarins have historically been implicated in plant defense against pathogens. Several studies have shown that coumarins accumulate upon pathogen infection and contribute to resistance by modulating redox homeostasis, affecting reactive oxygen species (ROS) dynamics and defense signaling (Beesley et al., 2023; Beyer et al., 2019; Stringlis *et al*., 2019; Weber Böhlen et al., 2026). These findings suggest that coumarins may serve dual functions in belowground and aboveground defense, linking rhizosphere interactions with systemic immune responses.

MYB72 induces *FERULOYL-CoA 6’-HYDROXYLASE1* (*F6’H1*), which encodes the key entry enzyme of simple coumarin biosynthesis in Arabidopsis that converts feruloyl-CoA into ‘6-hydroxyferuloyl-CoA (Kai et al., 2008; Zamioudis *et al*., 2014). This intermediate is subsequently converted into scopoletin by COUMARIN SYNTHASE (COSY) in roots (Vanholme et al., 2019). Scopoletin is reactive, and is stored *in planta* as its less reactive, glycosylated form scopolin in the vacuole (Robe et al., 2021b; Taguchi et al., 2000). It is remobilized through deglycosylation by BGLU42, enabling secretion into the rhizosphere (Stringlis *et al*., 2018b). Coumarin biosynthesis and secretion are strongly upregulated during Fe deficiency, and coumarin-deficient plants show impaired Fe uptake (Schmid et al., 2014). Under alkaline conditions, where Fe availability is limited by Fe(III)-OH precipitation, SCOPOLETIN 8 HYDROXYLASE (S8H) is induced in a MYB72-dependent manner to convert scopoletin into fraxetin (Siwinska et al., 2018), a coumarin that can solubilize Fe(III) from insoluble sources, thereby improving availability for plant uptake (Paffrath et al., 2024; Tsai et al., 2018). Although Fe deficiency can also trigger ISR-like phenotypes, these responses seem largely MYB72-independent and transcriptionally distinct from WCS417-induced ISR (Trapet et al., 2021; Zamioudis *et al*., 2014), indicating that under Fe-sufficient conditions rhizobacteria activate a partly overlapping but distinct metabolic program with that induced by Fe deficiency.

The shared regulation of WCS417-ISR and coumarin metabolism by MYB72 and BGLU42 suggests that coumarins contribute to the establishment of WCS417-ISR (Stassen et al., 2021; Zamioudis *et al*., 2014). Because coumarins can accumulate throughout the plant and potentially move between roots and shoots (Knox et al., 2018; Robe *et al*., 2021b), their spatial and temporal dynamics during ISR onset may provide insight into how rhizobacterial signals are translated into systemic immune responses. Here, we investigated the localization and dynamics of coumarin biosynthesis and accumulation across Arabidopsis roots and shoots in response to colonization of the roots by WCS417 using two-photon multispectral imaging, targeted metabolite profiling, root and shoot transcriptomics, and functional genetics. Our results revealed a coordinated activation of F6’H1-dependent coumarin biosynthesis along the root-shoot axis and demonstrate that coumarin accumulation is essential for WCS417-mediated ISR, supporting a role for MYB72–BGLU42-dependent coumarins as candidate signals linking rhizobacterial perception in roots to systemic immunity in leaves.

## RESULTS

### WCS417 induces spatially distinct coumarin accumulation patterns in roots

Root colonization by WCS417 activates ISR in Arabidopsis. This response requires activation of the root-specific transcription factor MYB72 and its downstream target BGLU42, which also regulate coumarin biosynthesis and activation in roots (Pieterse *et al*., 2021). Because coumarins can move systemically from roots to shoots (Robe *et al*., 2021b), they represent candidate signals linking rhizobacterial perception in roots to systemic immune responses in leaves. To investigate how WCS417 root colonization affects coumarin metabolism, we first examined coumarin accumulation directly in roots using two-photon multispectral imaging of coumarin autofluorescence. This approach enables *in situ* visualization of F6’H1-dependent coumarins, including scopolin and fraxin, at cellular resolution at the initial site of plant-bacteria interaction (Robe *et al*., 2021b). Following WCS417 treatment, the entire root system of Col-0 seedlings became rapidly colonized (Supplemental Figure S1). Enhanced autofluorescence of the coumarins scopolin (purple) and fraxin (green) was detected in primary and lateral roots of Col-0 seedlings grown for 5 days in the presence of WCS417 under Fe-sufficient conditions (50 µM Fe(III)-EDTA, pH 5.5; Figure 1A). Fraxin accumulation was consistently observed in the late maturation zone (LMZ) of the primary root. These observations are in accordance with previous reports showing that WCS417 induces the expression of coumarin biosynthesis genes even under Fe sufficient conditions (Zamioudis et al., 2015). F6’H1 catalyzes the committed entry step of the coumarin biosynthetic pathway and is a key determinant of coumarin accumulation in Arabidopsis (Kai *et al*., 2008). No fluorescence signal was observed in WCS417-treated roots of the coumarin biosynthesis mutant *f6’h1*, confirming the specificity of the detected coumarin signals (Figure 1A).

**Figure 1.**
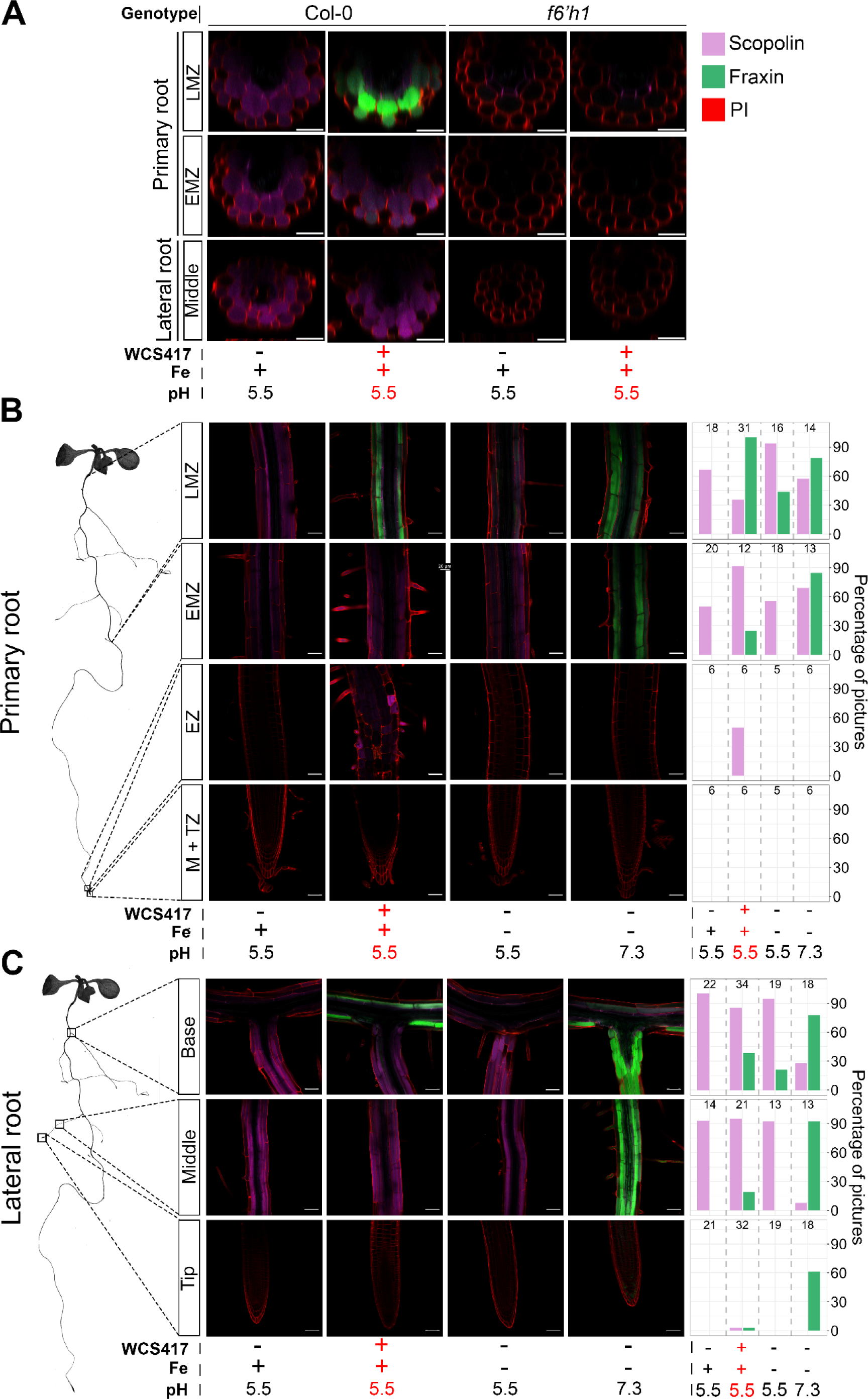
Merged two-photon, multispectral images of spatial patterns of WCS417- and Fe deficiency-induced coumarin accumulation in Arabidopsis Col-0 and *f6’h1* roots. **(A)** Detection of F6’H1-dependent coumarins in roots of 10-day-old Col-0 seedlings grown under Fe-sufficient conditions (+Fe, pH 5.5) following control or WCS417 treatments (See also Supplemental Fig. S2). For WCS417 treatment, 10 µL of a WCS417 suspension (OD₆₀₀ = 0.1 in 10 mM MgSO₄) was applied to the shoot-root junction at the time of transfer. After 5 days, seedlings were washed in sterile Milli-Q water, stained with propidium iodide (PI; 10 µg·mL^-1^, 1 min), and imaged by two-photon multispectral microscopy. Orthogonal views of Z-stacks obtained by two-photon multispectral imaging are shown for the early maturation zone (EMZ) and late mature zone (LMZ) of primary roots and for the middle part of lateral roots. No coumarin signal was detected in the coumarin biosynthesis mutant *f6’h1*, confirming specificity of the detected coumarin signals. Red fluorescence indicates PI-stained root cell walls. Seedlings were grown for 5 days on 1×MS medium and subsequently transferred to Hoagland medium containing +Fe (50 µM Fe(III)-EDTA, pH 5.5; Fe available), −Fe (0 µM Fe(III)-EDTA, pH 5.5; low Fe availability) or −Fe (0 µM Fe(III)-EDTA pH 7.3; very low Fe availability). **(B,C)** Coumarin accumulation in (**B**) primary and (**C**) lateral roots of Col-0 seedlings under the indicated WCS417, Fe, and pH treatments. Images are representative of three independent experiments, with 5-6 plants analyzed per treatment. The schematic plant (left) indicates imaging positions along the root axis, from the meristematic and transition zone (M+TZ), to the elongation zone (EZ), and the EMZ and LMZ. Frequency bars (right) indicate the percentage of images in which given coumarin signal was observed per treatment and location (numbers above bars denote total images analyzed). Representative images correspond to the most frequently observed coumarin patterns for each condition.

To compare WCS417-induced coumarin accumulation in Fe-sufficient plants with that triggered by Fe deficiency, roots of Col-0 seedlings were exposed for 5 days to WCS417 under Fe-sufficient conditions or two levels of Fe limitation: low Fe availability (0 µM Fe(III)-EDTA, pH 5.5), in which residual Fe from impurities in the agar and other medium components remains partly available, and very low Fe availability (0 µM Fe(III)-EDTA, pH 7.3), where alkaline conditions cause residual Fe to largely precipitate and become poorly accessible (Hsu et al., 2023) (Supplemental Figure S2). Two-photon multispectral imaging revealed that WCS417-treated roots grown under Fe-sufficient conditions displayed a distinct spatial pattern of coumarin accumulation along the primary root (Figure 1B), characterized by enhanced scopolin presence (purple) in the elongation zone (EZ) and early maturation zone (EMZ), and increased fraxin (green) in the EMZ and LMZ. These WCS417-induced patterns partly resembled those observed under Fe deficiency, particularly under very low Fe availability. However, coumarin accumulation patterns in lateral roots were more distinct (Figure 1C), with relatively higher fraxin abundance under very low Fe conditions than in WCS417-treated roots. Together, these results show that WCS417 induces a spatially defined and F6’H1-dependent coumarin signature in roots of plants growing under Fe-sufficient conditions. Although this response partially overlaps with canonical Fe-deficiency-induced coumarin patterns, it also displays treatment-specific features, indicating activation of a microbiota-associated coumarin program that diverges from classical Fe-deficiency responses.

### WCS417 induces local and systemic coumarin accumulation in roots and shoots

Coumarins such as scopoletin and its glycoside scopolin can also accumulate in Arabidopsis leaves, although typically at much lower levels than in roots (Kai et al., 2006; Robe *et al*., 2021b). To determine whether root colonization by WCS417 affects coumarin metabolism at the whole-plant level, we quantified coumarin accumulation in roots and shoots. Roots of 17-day-old *in vitro*-grown Col-0 seedlings were inoculated with WCS417, after which roots and shoots were harvested separately at 2, 5, or 7 dpi. Coumarin accumulation was then quantified using fluorescence detector-coupled HPLC (HPLC-FLD) analysis. Root colonization by WCS417 resulted in a pronounced increase in coumarin accumulation in roots (Figure 2A), confirming the two-photon multispectral imaging results (Figure 1). Levels of scopolin, fraxin, and scopoletin increased as early as 2 dpi, reaching up to 60-fold higher levels than in control plants, with fraxin showing the largest increase (Figure 2A).

**Figure 2.**
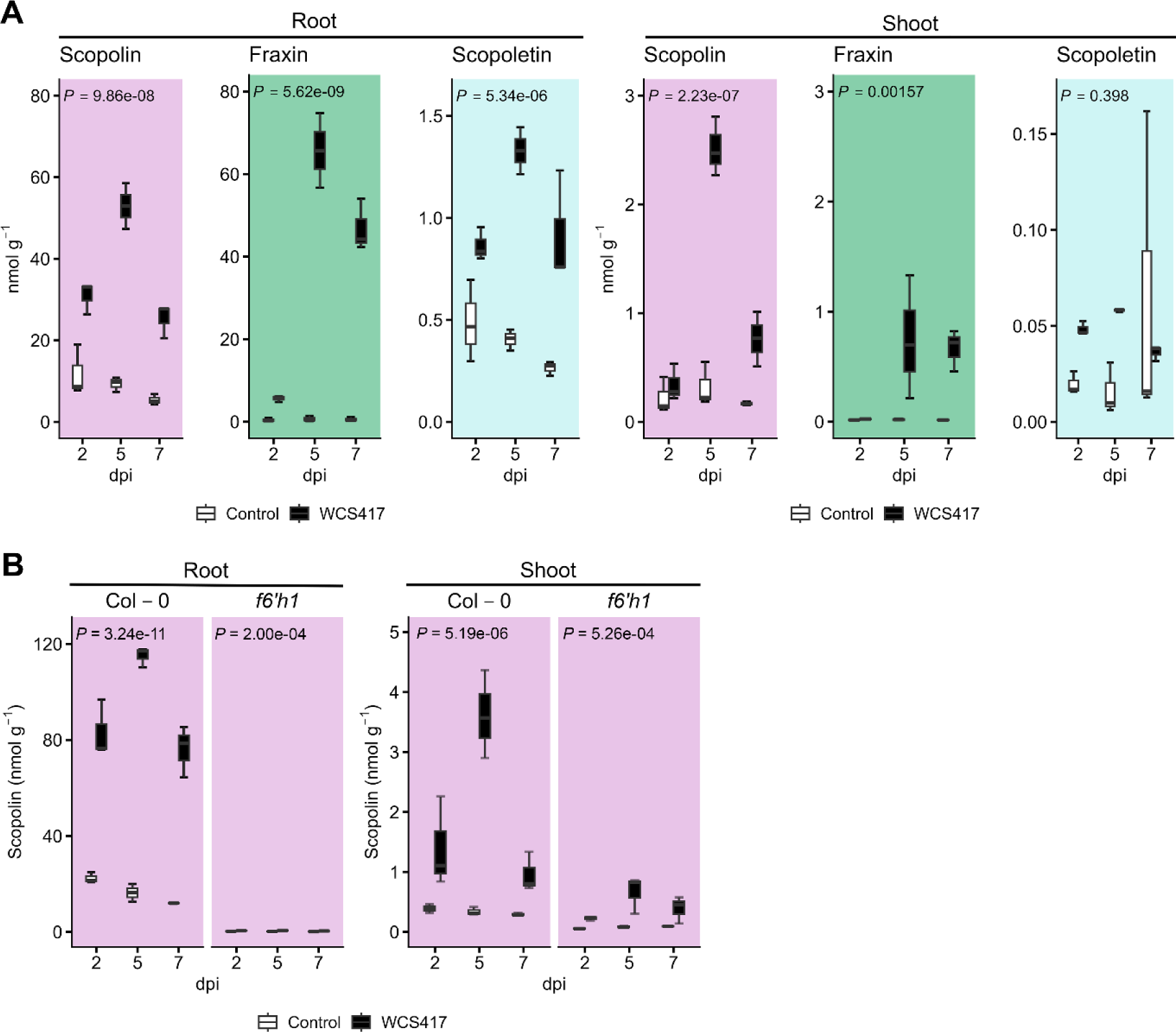
Coumarins level in roots and shoots of Arabidopsis Col-0 and f6’h1 upon WCS417 colonization. **(A)** HPLC-FLD quantification of scopolin (purple), fraxin (green), and scopoletin (blue) in roots and shoots of control and WCS417-treated Col-0 seedlings at 2, 5, and 7 dpi. *P* values indicate the significance of the treatment effect (Control vs. WCS417) based on two-way ANOVA (*n =* 2-3). **(B)** Scopolin accumulation in roots and shoots of Col-0 and the coumarin biosynthesis mutant *f6’h1* in control and WCS417-treated plants at the indicated time points. *P* values indicate significance of the treatment effect (Control vs. WCS417) based on two-way ANOVA (*n =* 3). Each sample consisted of 10-30 seedlings.

To investigate whether coumarins also accumulate in aerial tissues upon root colonization, we quantified coumarin levels in the shoot tissue. Levels of scopolin and fraxin were significantly higher in shoots of WCS417-treated plants than in control plants, and scopoletin followed a similar trend. However, their concentrations remained substantially lower than in roots (Figure 2A; note the different Y-axis scale between panels). In roots, WCS417-induced coumarin accumulation was F6’H1-dependent, as in the *f6’h1* mutant scopolin levels were hardly detectable (Figure 2B). In shoots, WCS417-induced scopolin levels were strongly reduced in *f6’h1* compared to Col-0, yet a small but significant increase in scopolin was still detectable in shoot tissue following WCS417 treatment of the roots (Figure 2B). This residual induction suggests that a minor F6’H1-independent route of coumarin biosynthesis may contribute to their accumulation. Whether the F6’H1 homolog F6’H2 or other enzymes are involved remains to be determined.

Together, these results demonstrate that root colonization by WCS417 triggers a systemic, F6’H1-dependent activation of coumarin biosynthesis, of which the strongest accumulation occurred in roots but with detectable increases also in shoots.

### WCS417 induces local and systemic transcriptional activation of coumarin biosynthesis genes

To determine whether the WCS417-induced coumarin accumulation observed at the metabolite level is supported by preceding transcriptional reprogramming, we performed time-resolved RNA-seq profiling of roots and shoots. Roots of 17-day-old *in vitro*-grown Col-0 seedlings were inoculated with WCS417, after which roots and shoots were harvested separately from 1 to 7 dpi. Analysis of all expressed genes (28,643), revealed that root colonization by WCS417 triggered pronounced transcriptional reprogramming in both tissues, with earlier responses in roots detectable from 1 dpi onwards (Figure 3A), where WCS417-treated samples already separated from controls (PERMANOVA: treatment R² = 0.34, *P* < 0.0001; timepoint R² = 0.29, *P* < 0.0001). In shoots, transcriptional changes emerged from 2 dpi onwards, when WCS417-treated samples began to diverge from controls (Figure 3A) (PERMANOVA: treatment R² = 0.16, *P* < 0.0001; timepoint R² = 0.49, *P* < 0.0001), consistent with root-to-shoot signaling.

**Figure 3.**
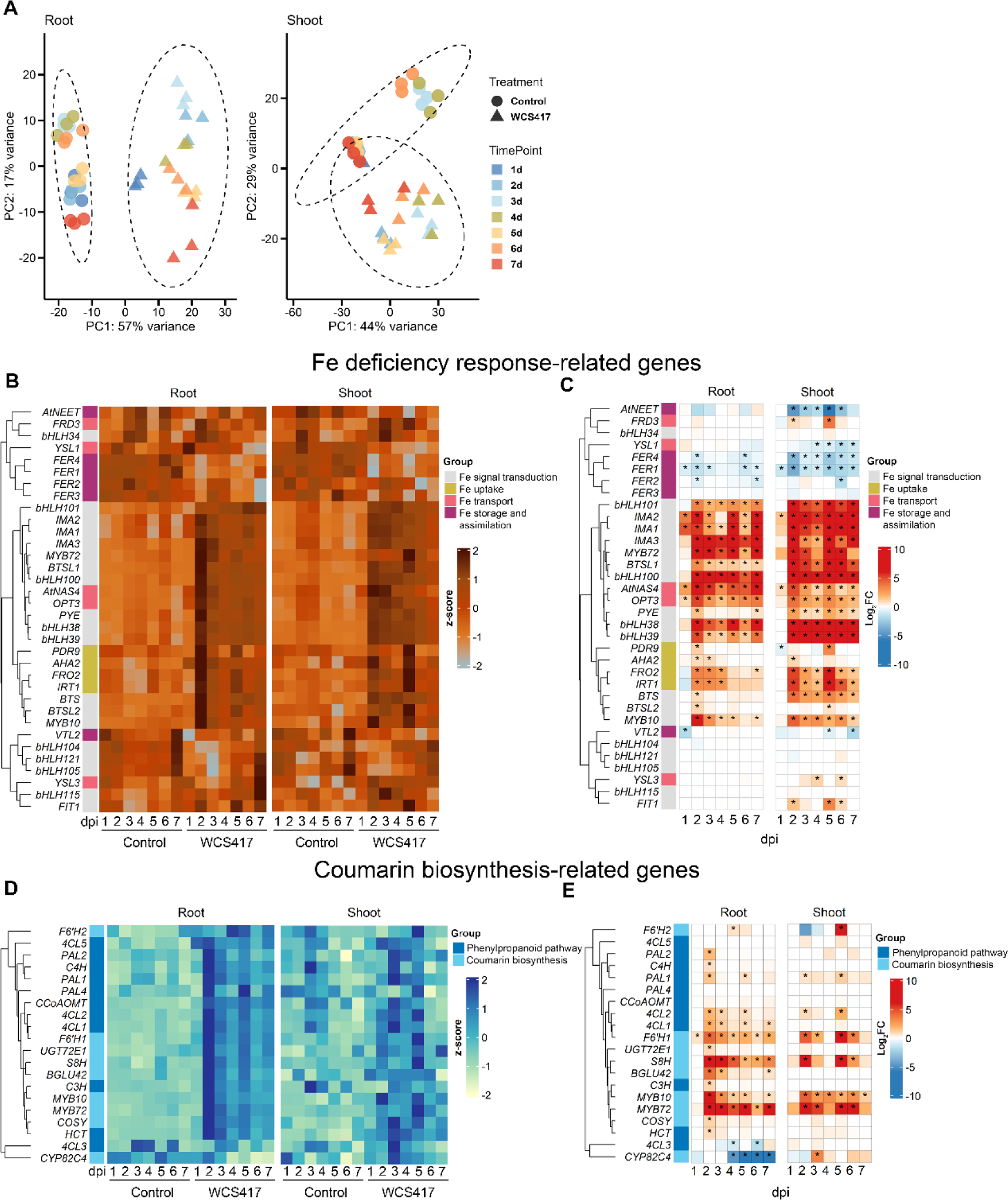
Plant-wide, time-resolved transcriptome reprogramming in response to root colonization by WCS417. **(A)** Principal component analysis (PCA) of transcriptome profiles of 28,643 genes in roots and shoots of control and WCS417-treated Arabidopsis Col-0 plants harvested at the indicated time points (1-7 dpi) for RNA-seq analysis. Each point represents one biological replicate. PC1, principal component 1; PC2, principal component 2. Three biological replicates per treatment and time point were included. **(B,D)** Heat maps showing z-scores of expression profiles of genes associated with (**B**) Fe-deficiency responses and (**D**) coumarin biosynthesis in roots and shoots following control or WCS417 treatment at 1–7 dpi. **(C,E)** Heat maps showing the corresponding WCS417-induced log₂ fold changes (log_2_FC) of the genes presented in panels (**B**) and (**D**). Gene expression values are based on normalized read counts. Color scales indicate relative expression levels from low to high (gray to dark red in panel **B** or yellow to blue in panel **C**) and log_2_FC values (**C,E**: red, upregulated; blue, downregulated following WCS417 treatment). Asterisks indicate significantly differentially expressed genes (WCS417 v.s. control, |log₂FC| > 1, *P*adj < 0.05). Genes are hierarchically clustered according to their expression patterns in roots.

Given the central role of coumarins in WCS417-mediated ISR and Fe acquisition, we focused specifically on a set of 34 Fe deficiency response-related genes and a set of 20 genes associated with the phenylpropanoid and coumarin biosynthetic pathways (Long et al., 2010; Mai et al., 2016; Robe *et al*., 2021a; Tissot et al., 2019; Wu et al., 2022; Zamioudis *et al*., 2014) (Supplemental Figures S3, S4, and S5; Supplemental Dataset D1). In roots, WCS417 strongly induced the expression of numerous Fe deficiency response genes (Figure 3B; z-scores), confirming previous findings (Zamioudis *et al*., 2014; Zamioudis *et al*., 2015). Analysis of the log_2_ fold changes (log_2_FC) (Figure 3C) revealed a coherent cluster of upregulated Fe response genes, including those involved in Fe deficiency signaling (e.g. *bHLH38/39/100/101/, IMA1/2/3, PYE, BTSL1/2, MYB10/72*), Fe uptake (*PDR9, AHA2, FRO2, IRT1*), and Fe transport (*e.g. AtNAS4, OPT3*), while all Fe storage-related genes were downregulated following WCS417 treatment. In parallel, 18 of the 20 examined phenylpropanoid and coumarin pathway genes were upregulated (Figures 3D and 3E), indicating coordinated root transcriptional activation of the pathway at the transcriptional level. The strongest induction was observed for key coumarin biosynthesis genes *MYB72, F6’H1,* and *S8H* consistent with activation of the canonical MYB72-dependent coumarin biosynthetic module.

Notably, Fe deficiency response and coumarin biosynthesis genes were also induced in shoots following root colonization by WCS417 (Figures 3B-E). Whereas transcripts of coumarin biosynthesis genes, including *MYB72*, *F6’H1*, and *S8H*, accumulated predominantly in roots, genes involved in systemic Fe homeostasis, such as the *IMA* genes and *OPT3,* showed higher transcript abundance in shoots (Supplemental Figures S4 and S5). Expression of coumarin biosynthesis genes peaked in roots at 2-3 dpi, whereas their induction in shoots reached a maximum at 5 dpi, mirroring the delayed accumulation of coumarins observed in shoots (Figure 2A). Together, these results demonstrate that WCS417 root colonization activates a coordinated and systemic transcriptional program for coumarin biosynthesis, characterized by rapid and strong induction in roots followed by delayed activation in shoots. This transcriptional reprogramming closely parallels the metabolite profiles and supports a model in which WCS417 triggers a F6’H1-dependent coumarin biosynthetic pathway operating locally in roots and systemically in aerial tissues.

### WCS417-ISR requires coumarin biosynthesis

Our imaging, metabolite, and transcriptomic analyses revealed that WCS417 root colonization induces a coordinated activation of coumarin biosynthesis along the root-shoot axis, resulting in strong local accumulation of coumarins in roots and a delayed but detectable increase in shoots. Because the ISR regulators MYB72 and BGLU42 control both coumarin biosynthesis and activation, these findings suggest that WCS417-induced coumarins may contribute functionally to the establishment of systemic resistance. If so, impairment of coumarin biosynthesis should compromise WCS417-induced ISR, whereas enhanced coumarin production should be sufficient to promote resistance.

To test this hypothesis, we performed ISR bioassays with 5-week-old soil-grown plants comparing Col-0 with the ISR-deficient *myb72* mutant and the coumarin biosynthesis mutant *f6’h1*. Across four independent experiments, WCS417 root colonization consistently induced ISR in Col-0 plants, as evidenced by a significant reduction in leaf disease incidence following challenge with the bacterial leaf pathogen *Pseudomonas syringae* pv. tomato DC3000 (*Pst*) compared with mock-treated plants (Figure 4A). As expected, the *myb72* mutant failed to express WCS417-induced ISR, confirming previous findings (Van der Ent *et al*., 2008; Zamioudis *et al*., 2014). Notably, the *f6’h1* mutant also failed to mount ISR, displaying similar disease levels in WCS417-treated and control plants. This loss of systemic resistance in *f6’h1* demonstrates that F6’H1-dependent coumarin biosynthesis is required for WCS417-induced ISR.

**Figure 4.**
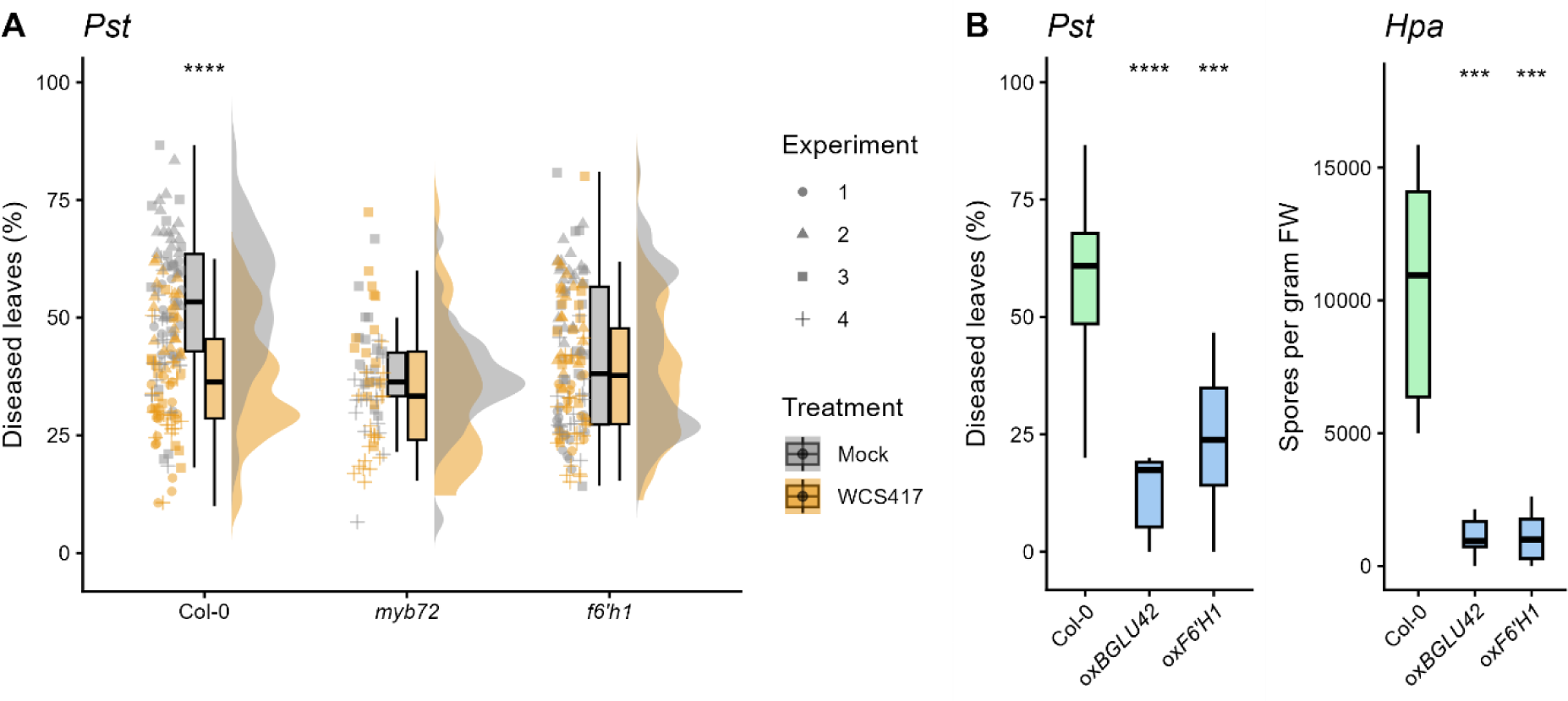
Effect of WCS417 and F6’H1 overexpression on disease resistance. **(A)** Disease incidence caused by *Pst* in 5-week-old Col-0, *myb72*, and *f6’h1* plants grown in soil with or without ISR-triggering WCS417 bacteria (5×10^7^ cfu·g^-1^ soil). Plants were co-cultivated with WCS417 in the root environment for 3 weeks prior to foliar infection with *Pst*. Disease incidence was scored as the percentage of leaves per plant with disease symptoms 4 days post infection. Statistical differences between treatments were assessed using a Wilcoxon rank-sum test on pooled data from four independent experiments. Points represent individual plants. Asterisks indicate significant differences between treatments (Wilcoxon rank-sum test; **** *P* ≤ 0.0001, each experiment contained *n* = 15-20 samples). **(B)** Disease incidence caused by *Pst* and spore counts of *Hpa* in 5-week-old mock-treated Col-0, ox*BGLU42*, and ox*F6’H1* plants, assessed 4 days post infection. Asterisks indicate significant differences compared with wildtype (Col-0) samples (Wilcoxon rank-sum test; \*\*\**P* ≤ 0.001, **** *P* ≤ 0.0001, *n =* 15-20).

To assess whether enhanced coumarin production is sufficient to promote systemic resistance, we next examined ox*F6’H1* plants overexpressing *F6’H1* (Schmid, 2014), which leads to enhanced accumulation of the coumarins scopoline and scopoletin in the shoots (Beesley *et al*., 2023; Weber Böhlen *et al*., 2026). In the absence of WCS417 treatment, ox*F6’H1* plants exhibited constitutively reduced disease incidence following infection with *Pst* (Figure 4B). Moreover, ox*F6’H1* plants displayed enhanced resistance to infection by the oomycete pathogen *Hyaloperonospora arabidopsidis* Noco2 (*Hpa*) (Figure 4B). The level of constitutively enhanced resistance to *Pst* and *Hpa* was comparable to that observed in the previously described ox*BGLU42* line (Zamioudis *et al*., 2014), suggesting that elevated coumarin levels confer broad-spectrum, ISR-like resistance against different types of pathogens.

Together, these findings provide functional validation that coumarin biosynthesis is both necessary and sufficient for the expression of WCS417-induced systemic resistance. In the context of the spatial, metabolic, and transcriptional activation of coumarin biosynthesis along the root-shoot axis described above, these results support a model in which MYB72-F6’H1-BGLU42-dependent coumarins act as key metabolic mediators linking rhizobacterial perception in roots to the establishment of systemic immunity in leaves.

### WCS417 modulates flg22-induced ROS production in a coumarin-dependent manner

Reactive oxygen species (ROS) production is one of the earliest cellular responses triggered by pathogen infection and plays a central role in immune signaling and defense activation. They have a central role during generation of the hypersensitive response in pattern- and effector-triggered immunity, and play a central role in modulation and propagation of many other plant stress responses. Because coumarins are redox-active metabolites with antioxidant properties that can modulate ROS homeostasis by scavenging excess ROS (Stringlis *et al*., 2019; Wu et al., 2023), we hypothesized that WCS417-induced systemic accumulation of coumarins may influence ROS dynamics during immune activation and thereby contribute to the establishment of ISR. To test this, we quantified the flg22-triggered oxidative burst in leaves of *in vitro*-grown seedlings using a luminol-based plate reader assay. Five-day-old plants were root-inoculated with WCS417 and cultivated for another five days as described above. Subsequently, the plants were rinsed, exposed to the bacterial MAMP flg22, and immediately assayed for ROS production.

As expected, flg22 treatment of control Col-0 plants triggered a rapid and transient ROS burst, characterized by a steep increase in luminescence shortly after elicitation (Figure 5A). In contrast, WCS417-treated Col-0 plants exhibited a significantly reduced ROS burst following flg22 treatment, indicating that root colonization by WCS417 significantly attenuates MAMP-induced ROS production in the shoot. This reduction in ROS accumulation is consistent with the increased levels of redox-active coumarins detected in WCS417-treated plants and suggests that these metabolites may contribute to quenching or buffering of the oxidative burst.

**Figure 5.**
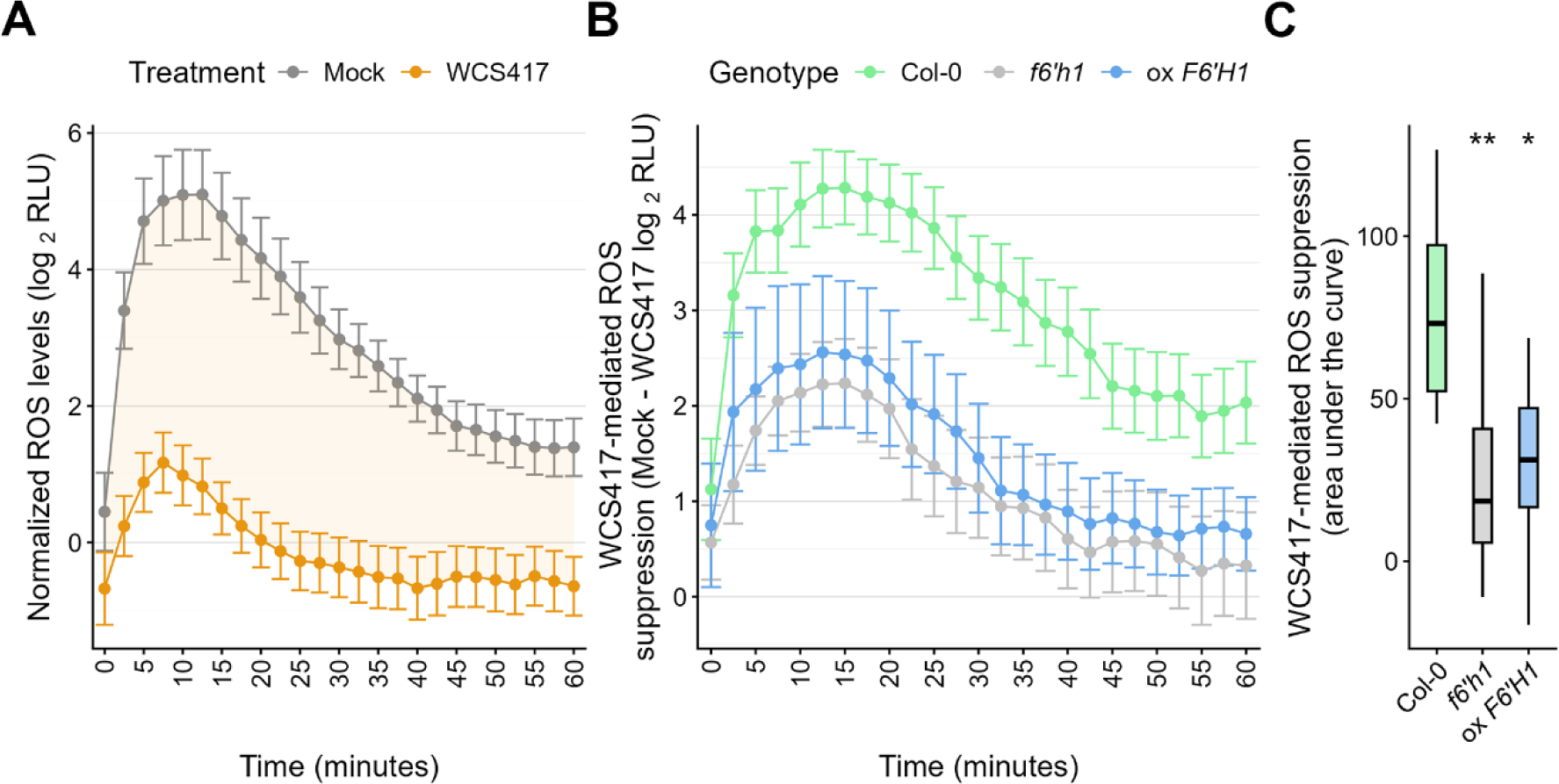
Effect of WCS417 root colonization on flg22-triggered ROS production in Col-0, f6’h1, and oxF6H1 plants. (A) Effect of WCS417 root colonization on flg22-triggered ROS production in leaves of *in vitro*-grown Arabidopsis Col-0 seedlings. ROS production was elicited in control and WCS417-treated seedlings with the bacterial MAMP flg22 or a Milli-Q mock treatment. ROS production was monitored for 60 min at 2.5-min intervals using a luminol-based assay, with relative luminescence units (RLUs) as output. Shown are log_2_-transformed RLU values of flg22-treated samples, normalized by subtracting the average of log_2_-transformed RLU values of their respective mock treatment. The highlighted area shows the difference in normalized ROS production between control and WCS417 pretreated plants. (B) Suppressive effect of WCS417 root colonization on flg22-triggered ROS production in leaves of Col-0, *f6’h1*, and ox*F6’H1* plants (the green Col-0 curve corresponds to the difference between mock and WCS417 in (**A**)). WCS417-mediated suppression of ROS production is expressed as the difference in normalized ROS production between control and WCS417-treated samples (± SE). (C) Quantification of WCS417-mediated suppression of normalized ROS production, expressed as the area under the curve (AUC) of the kinetics shown in (**B**). Asterisks indicate significant differences compared to control samples as determined by Welch’s *t*-test with Bonferroni-adjusted *P*-values (\**P*≤ 0.05; \*\**P* ≤ 0.01; *n =* 8).

Supporting this interpretation, the WCS417-mediated attenuation of the flg22-triggered ROS burst (calculated as ROS suppression in Figure 5B) was reduced in the coumarin-deficient *f6’h1* mutant (Figure 5B), in which coumarin levels are constitutively low. Accordingly, quantification of total ROS production (area under the curve; Figure 5C) showed that WCS417 treatment had a significantly weaker suppressive effect on the flg22-induced ROS burst in *f6’h1.* Notably, F6’H1-overexpressing (ox*F6’H1*) plants also showed a reduced WCS417-mediated suppressive effect on flg22-induced ROS production, possibly because WCS417 does not further enhance the already elevated coumarin levels in this line.

Together, these results indicate that WCS417-mediated activation of coumarin biosynthesis systemically affects ROS dynamics, modulating flg22-induced ROS accumulation in an F6’H1-dependent manner. By fine-tuning ROS responses while maintaining immune competence, coumarins may contribute to the primed defensive state characteristic of ISR, enabling beneficial rhizobacteria to enhance resistance without triggering detrimental overactivation of defense responses.

## DISCUSSION

### Coumarins as systemic mediators of WCS417-induced ISR

Rhizobacteria-mediated ISR has long been conceptualized as a root-localized perception event that primes distal tissues without leaving a major transcriptional footprint in shoots prior to pathogen challenge (Pieterse *et al*., 2014; Van Wees et al., 1999; Verhagen et al., 2004). However, the increased sensitivity and temporal resolution of RNA-seq now reveal a more dynamic scenario. Our time-resolved transcriptome analysis shows that WCS417 root colonization triggers rapid and extensive transcriptional reprogramming in roots from 1 dpi onwards, followed by a weaker but clearly detectable and temporally delayed transcriptional response in shoots starting at 2 dpi. This coordinated root-to-shoot wave parallels the sequential accumulation of coumarins in both tissues, indicating that ISR onset involves systemic metabolic reprogramming of coumarins rather than solely root-confined changes.

Mechanistically, we demonstrate that WCS417 activates a coordinated, F6’H1-dependent coumarin biosynthetic module along the root-shoot axis. Under Fe-sufficient conditions, WCS417 induces a spatially distinct coumarin signature in roots that only partially overlaps with canonical Fe-deficiency responses, highlighting activation of a microbiota-specific branch of the Fe-coumarin network. Strong local accumulation of scopolin, fraxin, and scopoletin in roots is followed by delayed but significant increases in shoots, accompanied by temporally aligned induction of *MYB72, F6’H1, S8H*, and related phenylpropanoid genes.

Together, these findings position coumarin biosynthesis as an active systemic component of ISR, linking rhizobacterial perception in roots to plant-wide transcriptional and metabolic reprogramming that accompanies the establishment of systemic immunity.

### Spatial regulation of coumarin accumulation in WCS417-colonized roots

Under Fe-sufficient conditions, genes involved in Fe uptake and coumarin biosynthesis are typically expressed at very low or undetectable levels, and coumarins, apart from a basal low level of scopolin, are generally not detected in Arabidopsis roots (Fourcroy et al., 2014; Robe *et al*., 2021b; Schmid *et al*., 2014; Zamioudis *et al*., 2014; Zamioudis *et al*., 2015). However, our gene expression profiling, imaging, and metabolite analyses demonstrate that colonization of the roots by WCS417 rhizobacteria activates the coumarin biosynthesis pathway, even in the absence of Fe deficiency. Importantly, activation of *MYB72* and other Fe-deficiency-responsive genes by WCS417 is not caused by bacterial competition for Fe. The main siderophore of WCS417, pyoverdine, has been shown to improve Arabidopsis Fe nutrition (Vansuyt et al., 2007), and both the volatile organic compounds of WCS417 as well as a siderophore-deficient WCS417 mutant were shown to induce the same MYB72-dependent Fe-deficiency and coumarin biosynthesis genes under Fe-sufficient conditions (Trapet *et al*., 2021; Zamioudis *et al*., 2015). This demonstrates that WCS417 activates this transcriptional and metabolic program through plant signaling rather than through physical Fe depletion by bacterial siderophores.

Using two-photon multispectral imaging, we provide new insights into the spatial organization of coumarin metabolism during root colonization by the ISR-inducing bacterium WCS417. WCS417 triggered distinct spatial patterns of coumarin accumulation along the longitudinal root axis: scopolin accumulated predominantly in younger root tissues, including the elongation zone and lateral roots, whereas fraxin was enriched in older tissues, particularly the late maturation zone. These distributions differ markedly from the spatial patterns reported for Fe-deficiency-induced coumarin accumulation (Robe *et al*., 2021b), suggesting that coumarin biosynthesis is locally regulated according to root developmental stage and/or microbial colonization. Interestingly, fraxetin accumulation is typically associated with alkaline conditions that trigger canonical Fe-deficiency responses (Robe *et al*., 2021b; Tsai *et al*., 2018), whereas WCS417 has been reported to acidify its surroundings through gluconic acid secretion (Yu et al., 2019). This apparent discrepancy may indicate that WCS417 activates MYB72-dependent coumarin biosynthesis independently of the canonical pH-dependent Fe-deficiency pathway, or alternatively that bacterial acidification varies spatially along the root axis. Such spatial specialization is consistent with the increasingly recognized view that plant-microbe interactions are highly compartmentalized along the root (Verbon et al., 2023). Indeed, recent studies have demonstrated that both root exudation and microbiome assembly are spatially structured, with distinct developmental root zones characterized by unique metabolite profiles and microbial communities (Galindo-Castañeda et al., 2024; Loo et al., 2024; Tsai et al., 2025). The WCS417-induced spatial coumarin signatures observed here may therefore likely reflect localized metabolic responses that shape microbial colonization and function along the root system. Together, these findings support the emerging concept that plants deploy specialized metabolites with high spatial precision, enabling local modulation of the rhizosphere microbiome during the establishment of ISR.

### Systemic coumarin accumulation: local synthesis or long-distance transport?

The detection of elevated coumarin levels in shoots following root colonization raises the question whether these metabolites are transported from roots or synthesized locally in aerial tissues. Although transcript levels of key biosynthetic genes were lower in shoots than in roots, their induction was clearly detectable and temporally aligned with shoot metabolite accumulation. This suggests that local biosynthesis and long-distance transport may both contribute.

Previous work has shown that coumarins such as scopolin and scopoletin can be detected in xylem sap and move from roots to shoots (Robe *et al*., 2021b). Conversely, the supplied coumarin glycoside of esculetin, esculin, was shown to travel from shoot to root through the phloem, indicating the possibility of complex coumarin trafficking in plant tissues (Knox *et al*., 2018). Our observation that scopoletin increased early in shoots supports the possibility of root-to-shoot transport. However, the residual WCS417-induced scopolin accumulation observed in *f6’h1* shoots suggests that a minor F6’H1-independent coumarin biosynthesis pathway may also operate in aerial tissues, possibly through F6’H2. Together, these findings indicate that both transport of root-derived coumarins to the shoot and local shoot biosynthesis of coumarins likely contribute to the WCS417-induced accumulation observed in aerial tissues. Determining the relative contribution of these two processes will require dedicated grafting experiments, ideally complemented by tissue-specific genetic complementation approaches.

### Coumarin biosynthesis is necessary and sufficient for ISR

Experiments with mutants *myb72* and *bglu42* already suggested that coumarin biosynthesis and metabolism are functional components of WCS417-induced ISR (Van der Ent *et al*., 2008; Zamioudis *et al*., 2014). Here, we show that the coumarin-deficient mutant *f6’h1* failed to mount WCS417-ISR, firmly establishing that coumarin production is indeed required for ISR. This is supported by our observation that *F6’H1* overexpression conferred constitutive resistance to both the bacterial pathogen *Pst* and the oomycete *Hpa*, to a degree comparable with *BGLU42* overexpression. These findings demonstrate that enhanced coumarin biosynthesis along the root-shoot axis is sufficient to induce broad-spectrum, ISR-like resistance. Importantly, the magnitude of shoot coumarin accumulation induced by WCS417 is modest compared to root levels, suggesting that even relatively small systemic increases can have biologically meaningful consequences for the onset of ISR. Coumarins are well-established phytoalexins and possess antimicrobial properties (Beesley *et al*., 2023; Beyer *et al*., 2019; Stringlis *et al*., 2019; Weber Böhlen *et al*., 2026), but their low concentrations in shoots under ISR conditions as reported here are unlikely to act solely via direct toxicity. Instead, their contribution to ISR may involve modulation of immune signaling processes.

### Coumarins fine-tune immune-associated ROS dynamics

ROS production is a hallmark of pattern-triggered immunity but must be tightly regulated, as excessive ROS can cause cellular damage and negatively impact growth. Coumarins such as scopoletin possess antioxidant properties and have been implicated in maintaining redox homeostasis (Stringlis *et al*., 2019). Here, we demonstrated that WCS417-mediated coumarin accumulation systemically modulates flg22-triggered ROS production in leaves in an F6’H1-dependent manner. In wild-type plants, root colonization by WCS417 significantly attenuated the MAMP-induced oxidative burst, whereas this attenuation was significantly reduced in the coumarin-deficient *f6’h1* mutant. Interestingly, the suppressive effect of WCS417 on flg22-triggered ROS production was also diminished in ox*F6’H1* plants. Given that these plants constitutively accumulate high levels of coumarins, this suggests that WCS417 does not exert an additional effect beyond an already elevated coumarin baseline.

Collectively, our data suggest that WCS417-induced systemic coumarin accumulation modulates ROS production without abolishing immune competence. Such modulation may contribute to the primed state characteristic of ISR, in which defense responses are optimized rather than constitutively activated.

### A feed-forward model linking rhizosphere interactions to systemic immunity

WCS417 is largely insensitive to the antimicrobial activity of coumarins and benefits from coumarin-mediated reshaping of the root microbiome (Stringlis *et al*., 2018b). This creates a potential feed-forward loop: WCS417 activates coumarin biosynthesis-related genes, such as *MYB72, F6’H1,* and *BGLU42*, enhancing coumarin biosynthesis and secretion; coumarins selectively suppress competing microbes while sparing WCS417; enhanced colonization reinforces ISR activation. Our data extend this model by demonstrating that coumarin biosynthesis also mediates systemic immune modulation. We propose a model in which rhizobacterial perception in roots activates a MYB72-F6’H1-BGLU42 module that drives spatially organized coumarin production (Fig. 6). Locally, coumarins contribute to Fe mobilization and microbiome assembly. Systemically, they accumulate in shoots where they fine-tune ROS dynamics and contribute to the establishment of an ISR-associated primed state. In this way, coumarins integrate nutritional, microbial, and immune signaling along the root-shoot axis. These findings strengthen the concept of the extended plant immune system (Pieterse, 2025), in which specialized metabolites operate as integrative signals coordinating belowground microbial interactions with aboveground immune competence.

**Fig. 6.**
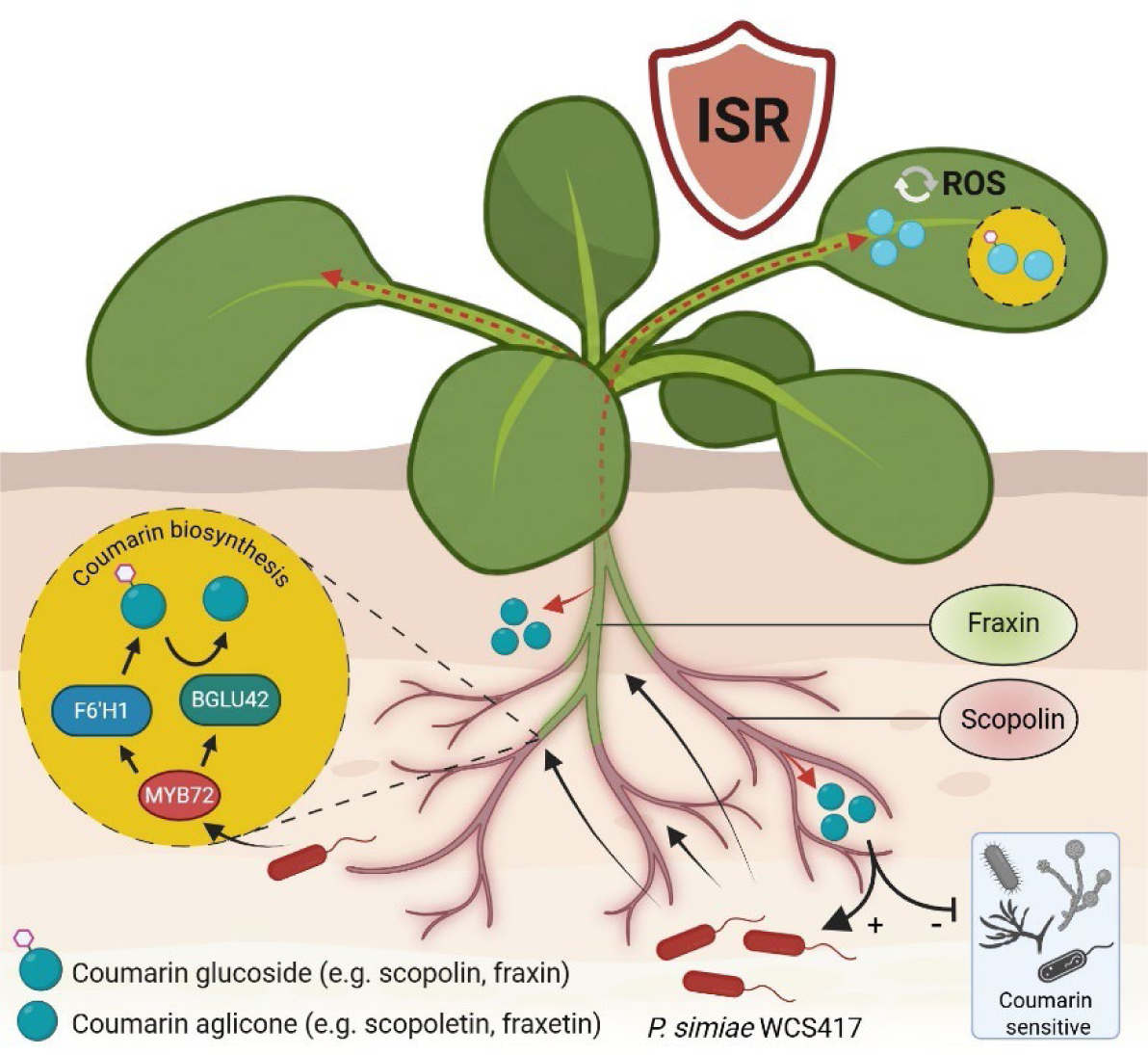
Model for the dual local and systemic functions of coumarins during WCS417-induced ISR. Root colonization by *Pseudomonas simiae* WCS417 activates the MYB72-F6’H1-BGLU42 coumarin pathway, resulting in enhanced coumarin biosynthesis and accumulation in roots. This response is spatially organized along the root axis, with distinct coumarin profiles associated with different developmental zones: e.g. scopolin accumulates predominantly in younger root tissues and lateral roots, whereas fraxin is more abundant in older root tissues, particularly the late maturation zone. Secreted coumarins can shape the root-associated microbial community through their selective antimicrobial activity: WCS417 is relatively tolerant to coumarins, whereas coumarin-sensitive microorganisms, including potential pathogens, are inhibited. WCS417 colonization also results in systemic activation of coumarin metabolism and increased coumarin accumulation in leaves. Shoot coumarins may originate from two, potentially complementary sources: transport of root-derived coumarins to the shoot and de novo coumarin biosynthesis in shoot tissues following systemic activation of the pathway. Genetic activation of coumarin metabolism through overexpression of either F6’H1 or BGLU42 is sufficient to phenocopy WCS417-induced systemic resistance, supporting a functional role for coumarin metabolism in ISR. In systemic leaves, increased coumarin accumulation is associated with modulation of ROS homeostasis and enhanced resistance against foliar pathogens. The identity of the coumarin(s) responsible for systemic resistance, their precise mode of action, and the relative contributions of root-derived transport and local shoot biosynthesis to the systemic coumarin pool remain to be determined. Image was created using Biorender.

## METHODS

### Plant material and growth conditions

*Arabidopsis thaliana* accession Col-0, mutants *f6’h1-1* (Kai *et al*., 2008) and *myb72-2* (Van der Ent *et al*., 2008), and overexpressing lines ox*F6’H1* 18A (Schmid, 2014) and ox*BGLU42* (Zamioudis *et al*., 2014) (all in Col-0 background) were used in this study. *In vitro*-grown Arabidopsis seedlings were cultivated as described by Hsu et al. (2023). In brief, seeds were surface-sterilized and sown on agar-solidified 1×Murashige and Skoog (MS) medium (Murashige and Skoog, 1962) supplemented with 4.7 mM MES, 0.5% sucrose, and 1% Plant Agar (Duchefa, Haarlem, the Netherlands). The medium pH was adjusted to 5.7 with 1 M KOH. Seeds were stratified for 48 h at 4 °C in complete darkness and plates were subsequently placed upright in a growth chamber. Seedlings were grown at 22 °C under short-day conditions (8 h light, 16 h dark; light intensity of 100 µM·m·s^-1^), and 70% relative humidity. Seedlings used for two-photon multispectral imaging were grown under similar conditions, except for long-day photoperiods (16 h light, 8 h dark). Prior to rhizobacterial inoculation of the roots, seedlings were transferred from germination plates to square Petri dishes (Greiner, Frickenhausen, Germany) containing solidified Hoagland medium (2 mM Ca(NO_3_)_2_, 5 mM KNO_3_, 2 mM MgSO_4_, 2.5 mM KH_2_PO_4_, 70 µM H_3_BO_3_, 14 µM MnCl_2_, 1 µM ZnSO_4_, 0.5 µM CuSO_4_, 10 µM NaCl, 0.2 µM, Na_2_MoO_4_), with or without 50 µM of Fe(III)-EDTA (treatment plates). Media were supplemented with 4.7 mM MES (for pH 5.5 plates) or HEPES (for pH 7.3 plates), 0.5% sucrose, and 1% Plant Agar, and the pH was adjusted to either 5.5 or 7.3 using 1 M KOH. After seedling transfer, plates were returned to the same growth chamber in upright position. Timelines and growth conditions used for the different experiments in this study are summarized in Supplementary Table S1.

### Cultivation of rhizobacteria and pathogens

Microbes used in this study included the ISR-inducing rhizobacterium *Pseudomonas simiae* WCS417 (WCS417) (Pieterse *et al*., 2021), an eGFP-labelled WCS417 strain constitutively expressing eGFP (eGFP-WCS417), the bacterial speck pathogen *Pseudomonas syringae* pv. *tomato* DC3000 (*Pst*) (Whalen et al., 1991), and the oomycete downy mildew pathogen *Hyaloperonospora arabidopsidis* Noco2 (*Hpa*) (Holub et al., 1994). The eGFP-WCS417 strain was generated by triparental mating between WCS417, *Escherichia coli* DH5α carrying the pMP4655 plasmid (Bloemberg et al., 2000), and *E. coli* HB101 harboring the helper plasmid pRK2073 (Better and Helinski, 1983).

WCS417 and *Pst* were stored at −80 °C in 25% glycerol containing 5 mM MgSO_4_. For cultivation, bacteria were streaked from glycerol stocks onto King’s medium B (KB) agar (King et al., 1954) plates supplemented with 50 µg·mL^-1^ rifampicin, and 10 µg·mL^-1^ tetracyclin when growing eGFP-WCS417, and incubated overnight at 28 °C. Bacterial cells were scraped from plates, resuspended in 10 mM MgSO_4_, after which 100 µL of the suspension was spread onto fresh KB plates with rifampicin for a second overnight incubation at 28 °C. Bacteria were then harvested, washed three times with 10 mM MgSO_4_ by centrifugation (5 min, 4,500 × *g*), and resuspended to an optical density at 600 nm (OD_600_) of 1.0 (10^9^ cfu·mL^-1^) or 0.1 (10^8^ cfu·mL^-1^), as required.

*Hpa* Noco2, an obligate biotrophic oomycete, was maintained by weekly propagation on Arabidopsis seedlings under conditions conducive to *Hpa* infection (16 °C, 10 h light/14 h dark, light intensity 100 μmol·m^−2^·s^−1^) as described previously (Goossens et al., 2023). Sporulating seedlings were harvested into 50-mL tubes containing 10 mM MgSO_4_ and vortexed vigorously to release spores. Spore density was determined using a hemocytometer and adjusted to 5 × 10^4^ sporangiospores mL^−1^ for inoculation.

### Rhizobacterial and pathogen inoculation and disease resistance assays

For *in vitro* assays, roots of Arabidopsis seedlings were inoculated with WCS417 by applying a 10-µL droplet of bacterial suspension (OD_600_ = 0.1) just below the root-shoot junction. Droplets were allowed to air-dry before plates were returned to the growth chamber and maintained in upright position (Stringlis et al., 2018a).

For disease resistance bioassays in soil, surface-sterilized Arabidopsis seeds were sown on twice-autoclaved river sand supplemented with liquid half-strength Hoagland medium (Van Wees et al., 2013). Seeds were stratified in darkness for 48 h at 4 °C and subsequently transferred to a short-day growth chamber (8 h light, 16 h dark, 22 °C, 70% relative humidity, light intensity 100 µM·m·s^-1^). After two weeks, uniform seedlings were transplanted into 60-mL pots containing a sterilized sand-soil mixture (5:12, v/v) either amended with WCS417 bacteria or 10 mM MgSO_4_ as a control. WCS417 was incorporated into the substrate prior to transplanting by thoroughly mixing a WCS417 suspension (OD_600_ = 1.0; 10^9^ cfu·mL^-1^) prepared in 10 mM MgSO_4_ into the sand-soil mixture to obtain a final density of 5 × 10^7^ cfu·g^-1^ soil. Control soils received an equal volume of 10 mM MgSO_4_. Plants were maintained under the same growth conditions throughout the experiment. For disease resistance assays, 15-20 biological replicates were used per treatment.

Three weeks after transplantation, the then 5-week-old plants were spray-inoculated with either *Pst* (OD_600_ = 0.1; 10^8^ cfu·mL^-1^) in 10 mM MgSO_4_ containing 0.015% (v/v) Silwet L-77 (Van Meeuwen Chemicals BV, Weesp, The Netherlands), or with *Hpa* (5 × 10^4^ sporangiospores mL^−1^) in sterile Milli-Q water as described previously (Goossens *et al*., 2023; Zamioudis *et al*., 2014). In the *Pst* pathosystem, disease symptoms were assessed 4 days post-inoculation by determining disease incidence (percentage of diseased leaves), as described previously (Van Wees *et al*., 2013). For *Hpa* infections, pathogen proliferation was quantified at day 4 after inoculation by spore counting as described previously (Goossens *et al*., 2023).

### Two-photon multispectral imaging

Images were acquired from 10-day-old, *in vitro*-grown seedlings that had been exposed to WCS417, low Fe, or control treatment for 5 days (Supplemental Figure S2). Imaging was performed as previously described (Robe *et al*., 2021b; Robe et al., 2023). Briefly, plants were removed from the agar plates, gently washed with sterile Milli-Q water, and stained with 10 μg·mL⁻¹ propidium iodide (PI) for 1 min prior to imaging to visualize the root cell walls.

Samples were imaged using an LSM 880 multiphoton microscope (Zeiss, Oberkochen, Germany) equipped with a Chameleon Ultra II laser (Coherent, Santa Clara, USA). Imaging relied on coumarin autofluorescence. Excitation was set at 720 nm (equivalent to 360 nm single-photon excitation), and emission signals were collected between 410 and 695 nm. Spectral imaging was performed using a GaAsP spectral detector with an 8.9 nm spectral resolution.

Mixed fluorescence signals from different compounds were separated on a pixel-by-pixel basis using the advanced linear unmixing function in Zen Black (Zeiss). Four predefined reference spectra were used to allow unmixing PI and the coumarins scopolin and fraxin (Robe *et al*., 2021b; Robe *et al*., 2023).

### Coumarin quantification by HPLC-FLD

Arabidopsis seedlings were transplanted to Hoagland medium at 7 days after germination and had been exposed to WCS417 or control treatment since day 17. Seedlings were co-cultivated with WCS417 bacteria for 2, 5 or 7 days prior to harvest. Coumarin detection was performed as described previously (Gao et al., 2020; Robe *et al*., 2021b).

Leaves and roots were harvested separately, washed in sterile Milli-Q water, flash frozen in liquid nitrogen, and ground for 1 min in the presence of glass beads using a TissueLyser II (Qiagen, Hilden, Germany) at 30 Hz. Coumarins were extracted in methanol:Milli-Q water (80:20, v/v), with solvent volumes adjusted to sample fresh weight to maintain an equal weight-to-solvent ratio across all samples within each experiment. Extracts were filtered through 0.45 µM filters (Sartorius AG, Göttingen, Germany), and equal volumes of filtrate were freeze-dried. Freeze-dried samples were resuspended in 100 µL of Milli-Q water containing 10% methanol, 9% acetonitrile and 0.09% formic acid.

Samples were analyzed by high-performance liquid chromatography (HPLC) using the 1220 Infinity II LC system (Agilent Technologies, Santa Clara, USA) coupled to a fluorescence detector Prostar 363 fluorescence detector (Varian, Palo Alto, USA) as described previously (Gao *et al*., 2020; Robe *et al*., 2021b). Coumarin concentrations were calculated in nmol·g^-1^ fresh weight based on the evaporated volume, total extraction solvent volume, and compound-specific response factors.

### RNA isolation and sequencing

Plant material was harvested from seedlings that were transplanted to Hoagland medium at 7 days after germination and subsequently exposed at day 17 to WCS417 or a control treatment for 1 to 7 days. For RNA sequencing analysis of plant transcriptional profiles, samples were collected 1 to 7 days post-inoculation (dpi). Shoots and roots were carefully separated, rinsed in sterile Milli-Q water, and flash-frozen in liquid nitrogen. Samples were stored at −80 °C until further processing. Each treatment and time point consisted of three biological replicates, with each replicate comprising pooled material from thirty root systems or eight shoots harvested from similarly treated plants.

RNA isolation and library preparation were performed as described by Stringlis et al. (2018a). Briefly, total RNA was extracted using the RNeasy Plant Mini Kit (Qiagen) according to the manufacturer’s instructions. Residual genomic DNA was removed by on-column DNase digestion using the RNase-Free DNase Set (Qiagen). RNA integrity and quality were assessed using an Agilent Bioanalyzer with the Agilent RNA 6000 Nano Kit (Agilent Technologies). Samples with an RNA integrity number (RIN) ≥ 8 were selected for library preparation.

Strand-specific libraries were generated using the NEBNext Ultra II Directional RNA Kit (New England Biolabs, Ipswich, USA), in combination with the NEBNext poly(A) mRNA Magnetic Isolation Module. Libraries were sequenced on an Illumina NextSeq platform (Illumina, San Diego, USA) in paired-end mode (2 × 150 bp), yielding approximately 40 million reads per sample. Raw RNA-seq read data have been deposited in the European Nucleotide Archive (ENA) under project accession number PRJEB122822.

Reads were pseudoaligned to the TIR10 cDNA reference (Lamesch et al., 2012) using Kallisto (v0.45.0) with 100 bootstrap iterations and default parameters (Bray et al., 2016). Transcript-level abundances were summarized to gene-level counts using tximport (v1.26.1) (Soneson et al., 2015). Differential gene expression analysis was performed using DESeq2 (v1.38.3) (Love et al., 2014), including normalization of raw counts, variance-stabilizing transformation (VST), and identification of differential expressed genes relative to the corresponding control treatment.

### Measurement of flg22-induced reactive oxygen species (ROS) production

ROS production was quantified in flg22-treated Arabidopsis plants using a luminol-based assay adapted from a previously described method for flg22-triggered oxidative burst measurement (Bisceglia et al., 2015). In brief, 10-day-old *in vitro*-grown plants that had been exposed to WCS417 or control treatment for 2 days as described above were separated at the root-shoot junction, and 2-3 shoots were placed in 96-well plates, immersed in 150 µL of Milli-Q water. Milli-Q water was refreshed 4 times during 2-3 hours, after which the plates were covered with aluminum foil overnight and placed in a long-day growth chamber (described earlier) to eliminate ROS production caused by cutting. The Milli-Q was carefully removed and replaced with 250 µL reaction mixture (30 µM luminol L-012; Sigma-Aldrich, Saint Louis, USA) and 5 × 10^4^ U µL horseradish peroxidase (HRP; Sigma-Aldrich) diluted in Milli-Q water and 0.5 µM flg22 (GenScript Biotech, Nanking, China) or the same volume of Milli-Q as control. Luminescence in each well was recorded on a Glomax luminometer (Promega, Madison, USA) at 2.5-min intervals. Log_2_-transformed relative luminescence units were used for subsequent data analysis. Means of the Milli-Q treated samples were subtracted from flg22-treated samples per treatment as normalization.

### Fluorescence imaging of eGFP labelled-WCS417

eGFP-WCS417 was used to monitor bacterial colonization dynamics on Arabidopsis roots over a period of seven days. Col-0 seedlings were grown and treated as described for the RNA-seq experiment. Fluorescence images were acquired once every 12 h using the HADES root phenotyping system at the Netherlands Plant Eco-phenotyping Centre (NPEC) (Pereira-Mendes et al., 2026). eGFP fluorescence was detected using an excitation wavelength of 470 nm and a filter set with an emission window of 513-520 nm. Imaging parameters were set to a shutter speed of 1000 ms and a detector sensitivity of 1% (FluorCam2 imaging unit Photo Systems Instruments (PSI), Drásov, Czech Republic).

### Software

All data processing, visualization, and statistical analyses were performed in R (R Core Team, version 4.4.2) using the packages ComplexHeatmap (v2.25.3), cowplot (v1.2.0), DESeq2 (v1.38.3), dplyr (v1.1.4), ggdist (v3.3.3), gghalves, ggplot2 (v4.0.0), ggpubr (v0.6.1), ggtext (v0.1.2), pracma (v2.4.4), rstatix (v0.7.2), tidyr (v1.3.1) and tidyverse (2.0.0).

## Supporting information

Supplemental DatasetD1_HsuStassen

## AUTHOR CONTRIBUTIONS

Conceptualization, S.-H.H., M.J.J.S., K.R., C.D., I.A.S., and C.M.J.P.; methodology, S.-H.H., M.J.J.S., K.R., and E.I.; formal analysis, S.-H.H., and M.J.J.S.; investigation, S.-H.H., M.J.J.S., K.R., E.I., C.D., C.M.J.P., and I.A.S.; writing - original draft, S.-H.H., and M.J.J.S.; writing - review & editing, S.-H.H., M.J.J.S., K.R., E.I., C.D., I.A.S., and C.M.J.P.; visualization, S.-H.H., M.J.J.S., I.A.S., and C.M.J.P.; funding acquisition, C.M.J.P. and C.D.

## ACKNOWLEDGMENTS

The authors thank Ricardo Giehl for kindly providing the ox*F6’H1* seeds.

## DATA AVAILABILITY

Raw RNA-seq read data have been deposited in the European Nucleotide Archive (ENA) under project accession number PRJEB122822.

## DECLARATION OF GENERATIVE AI AND AI-ASSISTED TECHNOLOGIES IN THE WRITING PROCESS

During the preparation of this work the authors used ChatGPT in order to improve readability and language. After using this tool/service, the authors reviewed and edited the content as needed and take full responsibility for the content of the publication.

## SUPPLEMENTAL INFORMATION

**Supplemental Figure S1.**
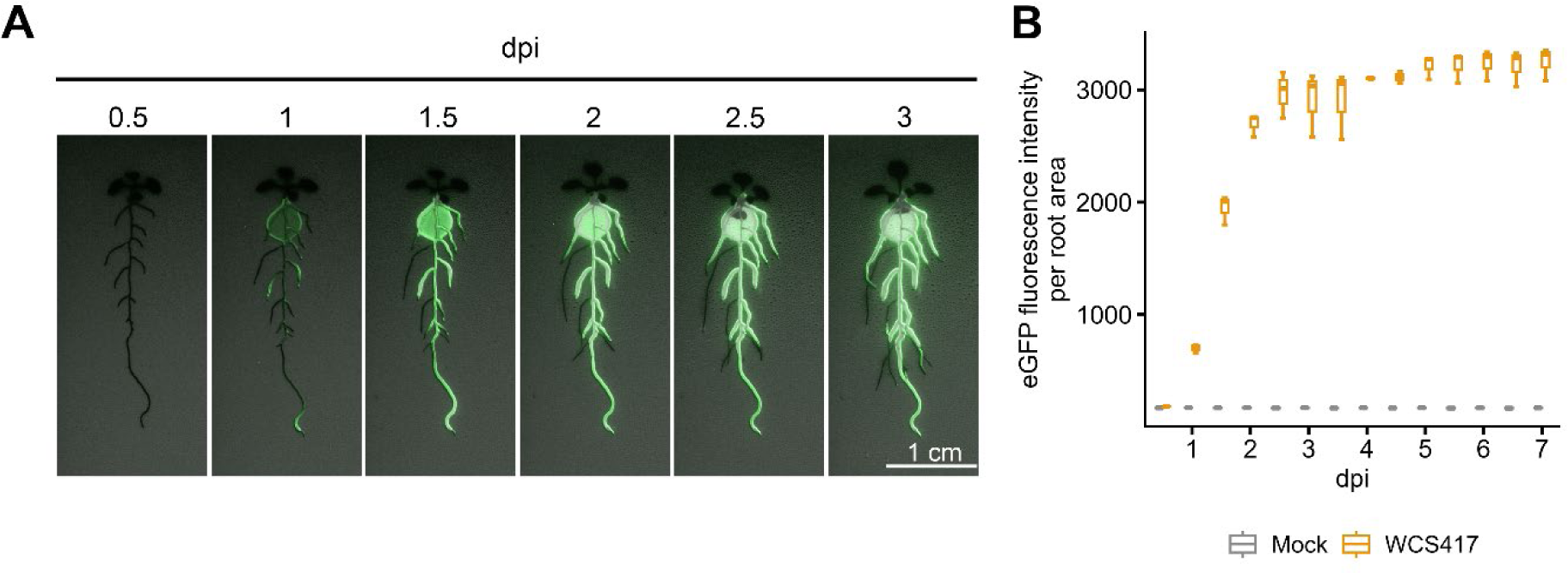
Time course of WCS417 colonization of Arabidopsis roots. Representative images of Arabidopsis Col-0 seedlings grown on agar plates and inoculated with eGFP-labeled WCS417 at the root-shoot junction. **(A)** Seedlings are shown at 0.5 to 3 dpi, illustrating the progression of bacterial colonization along the root system over time. eGFP fluorescence indicates the presence of WCS417 on the root surface and surrounding rhizosphere. Images are representative of multiple independent experiments showing similar colonization dynamics. **(B)** Quantification of bacterial fluorescence per root area (pixel). Data represent three biological replicates per time point, each consist of five seedlings.

**Supplemental Figure S2.**
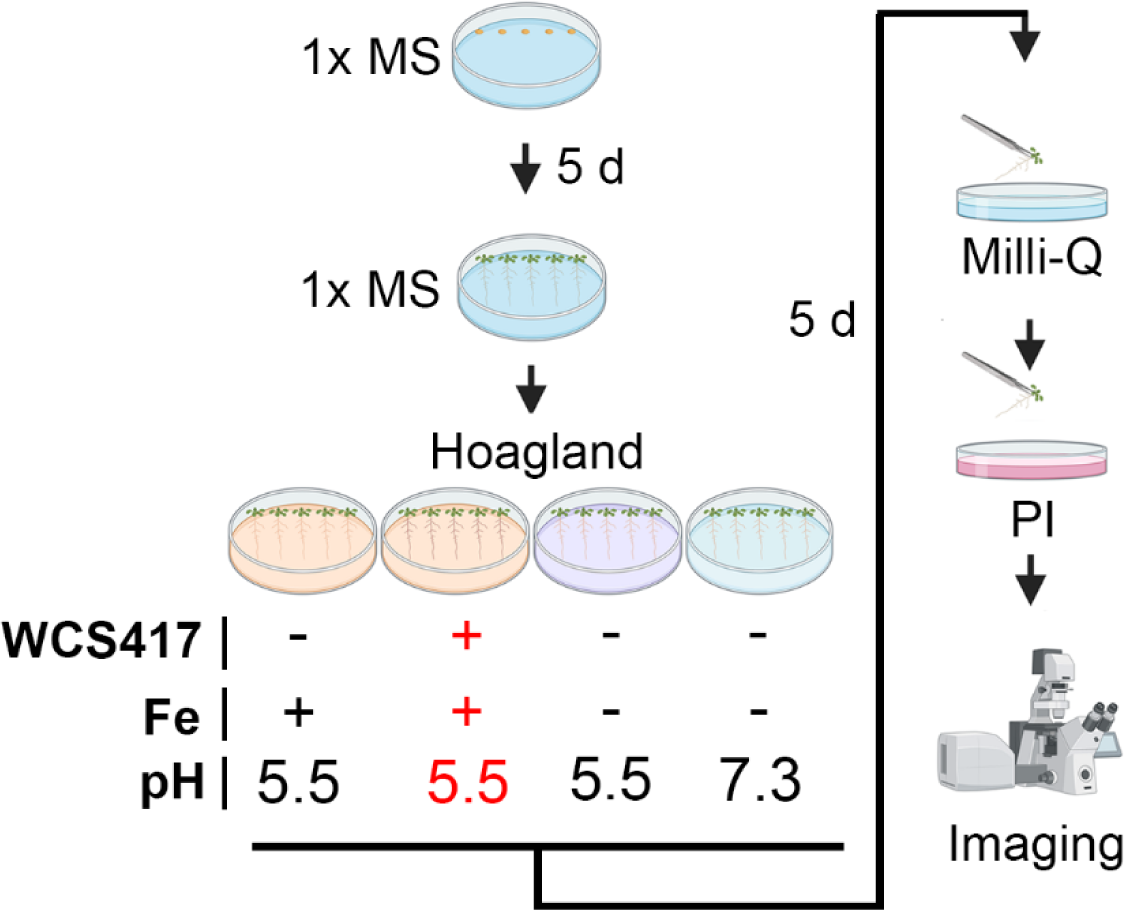
Experimental setup used for comparative visualization of coumarin accumulation under Fe deficiency and WCS417 treatment. Schematic overview of procedures used to visualize coumarins under Fe deficiency and WCS417 treatment conditions. Five-day-old Arabidopsis seedlings were transferred from 1× MS to agar-solidified Hoagland medium containing either sufficient Fe (+Fe, pH 5.5), low Fe (-Fe, pH 5.5), or very low Fe availability (-Fe, pH 7.3). WCS417 was applied to the roots at the time of transfer to the treatment plates. After 5 days, Arabidopsis plants were removed from the agar plates, washed with sterile Milli-Q water, and stained with 10 μg·mL⁻¹ propidium iodide (PI) prior to imaging to visualize the root cell walls. Images were acquired after 10 days of total seedling growth, across 3 experiments, with 5-6 plants per treatment. Image was created using Biorender.

**Supplemental Figure S3.**
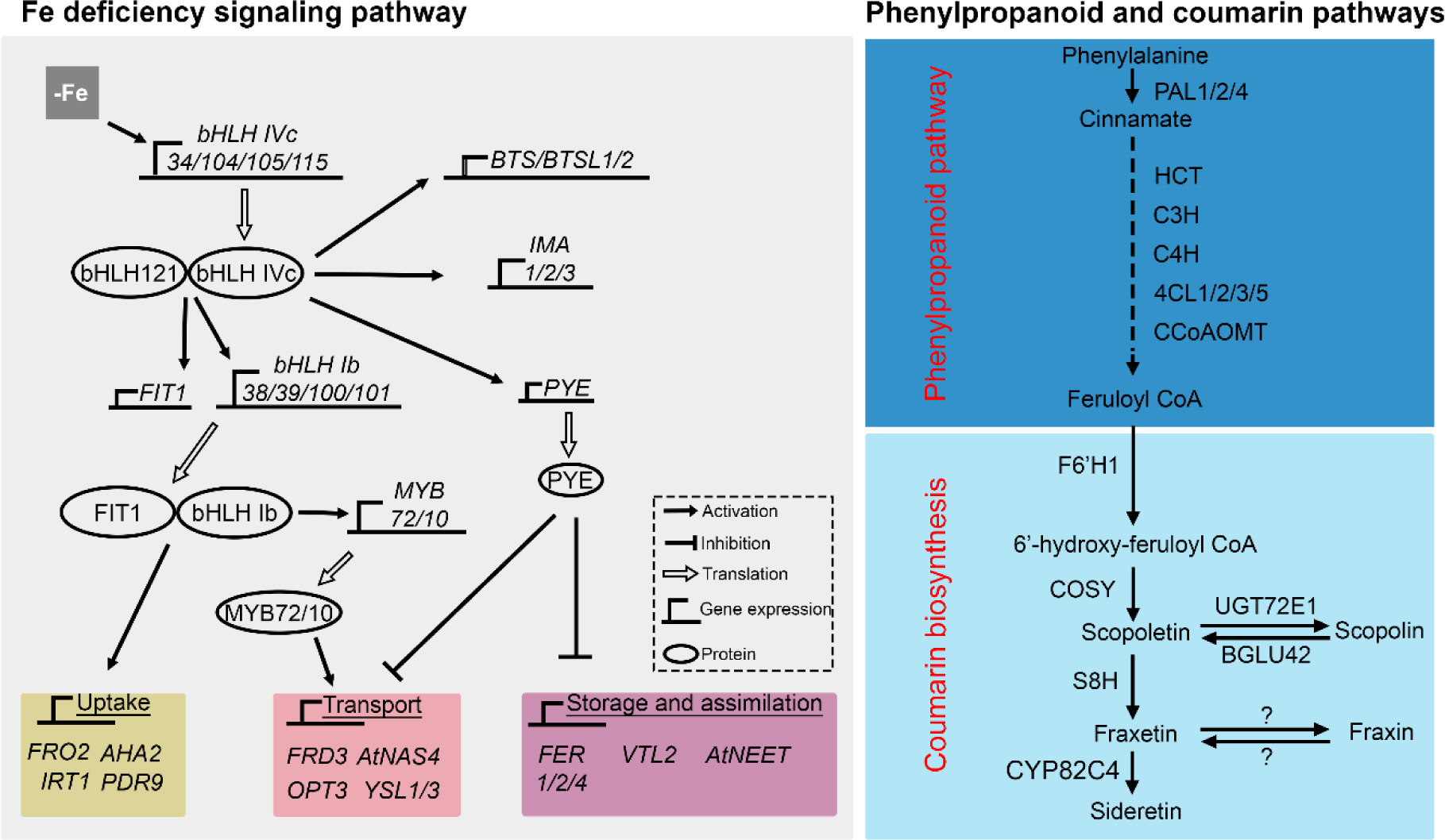
Schematic overview of the Fe-deficiency signaling network and the phenylpropanoid–coumarin biosynthesis pathway. Simplified representation of the molecular network underlying the Fe-deficiency response, including key transcription factors, regulatory components, and the machinery involved in Fe uptake, transport, storage and assimilation (left). The right panels depict the major steps of the phenylpropanoid pathway leading to the synthesis of feruloyl-CoA, the precursor of simple coumarins, and the downstream F6’H1-dependent biosynthesis and metabolism of coumarins, including scopoletin, scopolin, fraxetin, and fraxin. Many of the components shown correspond to genes that were significantly differentially expressed in response to WCS417 treatment in the transcriptome analysis presented in this study. Pathway components were compiled from studies on Fe deficiency signaling and phenylpropanoid/coumarin biosynthesis (Long *et al*., 2010; Mai *et al*., 2016; Robe *et al*., 2021a; Tissot *et al*., 2019; Wu *et al*., 2022; Zamioudis *et al*., 2014).

**Supplemental Figure S4.**
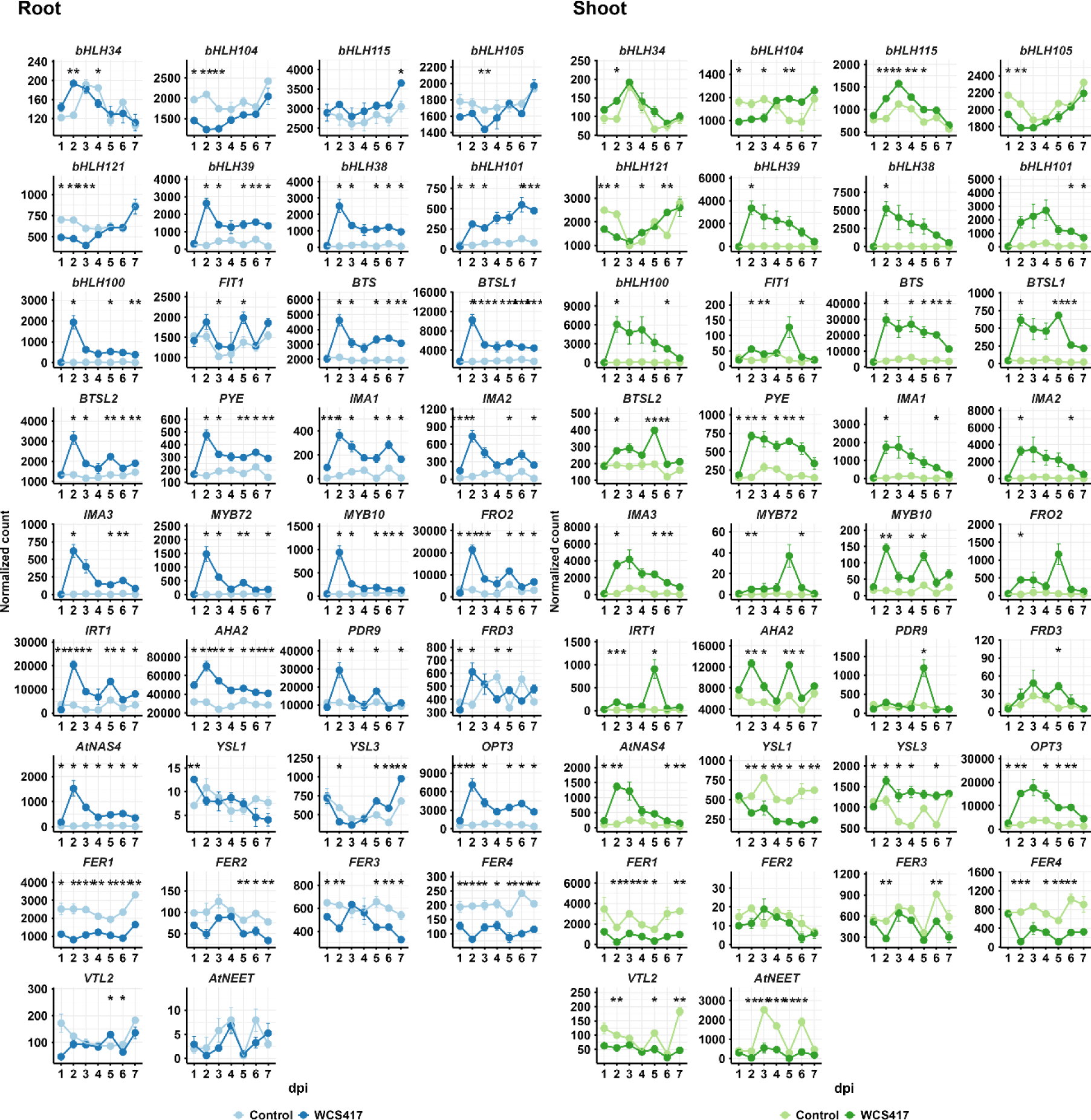
Expression patterns of 34 Fe deficiency response-related genes in roots and shoots of control and WCS417-treated Arabidopsis Col-0 plants. Time-course expression profiles of selected Fe-deficiency response genes extracted from the RNA-seq dataset. Expression levels are presented as normalized read counts. Roots of 17-day-old seedlings were inoculated with WCS417, and root and shoot samples were collected at 1, 2, 3, 4, 5, 6, and 7 dpi. Values represent means ± SE of three biological replicates. Asterisks indicate statistically significant differences compared with control samples (Welch’s *t* test; \**P* < 0.05, \*\**P* < 0.01, \*\*\**P* < 0.001).

**Supplemental Figure S5.**
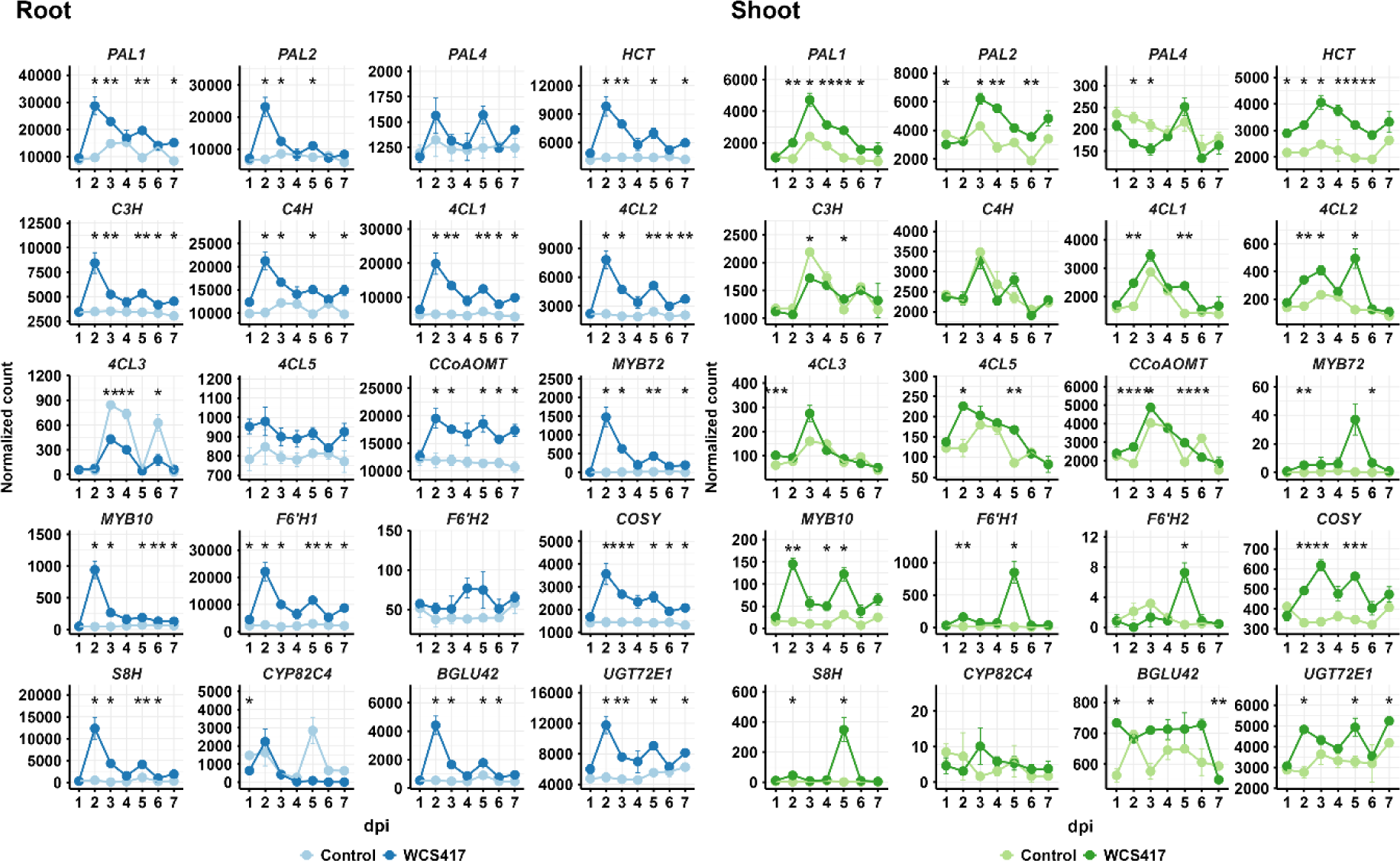
Expression patterns of phenylpropanoid and coumarin biosynthesis-related genes in roots and shoots of control and WCS417-treated Arabidopsis Col-0 plants. Time-course expression profiles of 20 selected phenylpropanoid and coumarin biosynthesis-related genes extracted from the RNA-seq dataset. Expression levels are presented as normalized read counts. Roots of 17-day-old seedlings were inoculated with WCS417, and root and shoot samples were collected at 1, 2, 3, 4, 5, 6, and 7 dpi. Values represent means ± SE of three biological replicates. Asterisks indicate statistically significant differences compared with control samples (Welch’s *t* test; \**P* < 0.05, \*\**P* < 0.01, \*\*\**P* < 0.001).

**Supplemental Table S1.** Experimental timelines and growth conditions for Arabidopsis assays. Overview of sampling schemes used for transcriptomics, metabolite profiling, imaging, ISR assays, and ROS measurements. Time points indicate days (d) or weeks (w) after sowing. The Table summarizes the timing of seedling transfer to treatment plates (Transplanting), application of *Pseudomonas simiae* WCS417 to the roots (WCS417 treatment), pathogen inoculation or flg22 treatment, and tissue harvesting and disease scoring for each experiment. Time points are indicated as days (d) or weeks (w) after sowing. Growth conditions refer to photoperiod regimes used throughout each experiment (short-day: 8 h light/16 h dark; long-day: 16 h light/8 h dark).

| Experiment | Transplanting | WCS417 treatment | Pathogen/<br>flg22 treatment | Harvesting /<br>scoring | Growth conditions |
| --- | --- | --- | --- | --- | --- |
| RNA-seq | 7 d | 17 d | – | 18, 19, 20, 21, 22, 23, 24 d | Short-day |
| Two-photon<br>multispectral imaging | 5 d | 5 d | – | 10 d | Long-day |
| HPLC-FLD | 7 d | 17 d | – | 19, 22, 24 d | Short-day |
| ISR bioassay | 2 w | 2 w | 5 w (*Pst*, *Hpa*) | 5 w + 4 d | Short-day |
| ROS assay | 5 d | 7 d | 10 d (flg22) | every 2.5 min for 62.5 min | Short-day |

**Supplemental Dataset D1. Gene expression profiles of 34 Fe deficiency response-related genes and 20 phenylpropanoid and coumarin biosynthesis-related genes in Arabidopsis roots and shoots in response to root colonization by P. simiae WCS417 from 1 to 7 dpi. See separate file.**

